# History of recurrent short and long-term coral bleaching events in Indian coral reefs: a systematic review of contrasting bleaching patterns, lessons learned, and future directions

**DOI:** 10.1101/2024.05.18.594244

**Authors:** Thinesh Thangadurai, Kalyan De, Sobana Murugesan, P Sivagurunathan, Riana Peter, Pasiyappazham Ramasamy, Joseph Selvin, Polpass Arul Jose, Anthony Bellantuono

## Abstract

Climate change has intensified coral bleaching events, leading to widespread coral mortality and threatening the coral reef survival worldwide. However, species susceptibility patterns to bleaching differ spatially and temporally, making it crucial to understand bleaching impact and species susceptibility patterns across latitudinal ranges, particularly from understudied locations. We critically reviewed 23 years of bleaching episodes on Indian coral reefs (four major reefs and other patch reefs) reported from 1998 to 2020 to understand the geographical footprint of bleaching patterns, species susceptibility differences, and their impact. We found that all four major Indian reef systems (Gulf of Kachchh, Gulf of Mannar, Lakshadweep, and the Andaman Islands) have experienced three major bleaching episodes (1998, 2010, and 2016) and multiple short-term bleaching events. Short-term bleaching events (DHWs> four weeks) created differences in species susceptibility, but the disparity among species diminished to long-term bleaching events (DHWs <4 weeks) in all reef sites. *Acropora*, *Porites*, *Pocillopora*, *Montipora,* and *Turninaria* in the Gulf of Mannar, *Porites,* and *Favia* in the Gulf of Kachchh, and *Acropora* in the Andaman and Lakshadweep were the most vulnerable genera during all the bleaching events. In terms of recovery, *Porites* and other massive forms recovered better than branching forms. Furthermore, species susceptibility pattern analysis (winners vs. losers) to 2016 long-term bleaching events found conflicting results from studies (one-time surveys) in the same area. This result indicates that a one-time survey (survey timing) influences results as bleaching events lengthen, necessitating repeated surveys to determine winners and losers. Although inconsistent data from different study sites makes it difficult to create predictive models, data collected over the past 23 years provides critical insight into coral community susceptibility patterns and bleaching impacts, underscoring the need for systematic, uniform surveys in the future.

## Introduction

Increasing coral bleaching frequency threatens the persistence of coral reef ecosystems, which provide livelihoods, food security, and coastal protection to over 280 million people worldwide (Moberg and Folke, 1999, Hughes et al., 2018). Coral bleaching is a multifaceted environmental problem that can arise due to various factors, such as increased seawater temperatures, extremely low or cold temperatures, salinity depletion, pollution, ocean acidification, and excessive exposure to sunlight (Hughes et al., 2018). However, the primary cause of coral bleaching is identified as increased episodes of marine heatwaves due to ongoing climate change. These climate-induced coral bleaching events occur when reef-building corals expose to a few degrees above the average summer temperature, causing a break down in the symbiotic relationship between coral and their endosymbiotic algal partners called *Symbiodinacea* (LaJeunesse et al., 2018, Hughes et al., 2018). This symbiotic relationship between corals and their intracellular algal symbiosis is the engine of the reef that forms the basis of its highly diverse ecosystem structure. Bleached autotrophic corals are physiologically deprived of regular food supply in the absence of their algal partner (Davy et al., 2012), and prolonged bleaching events led to extensive coral die-off (Glynn et al., 2001; González-Pech et al., 2021; Hughes et al., 2017b). Unfortunately, coral bleaching episodes have become more frequent and severe as the number of warmest years has increased, with nine of the ten warmest years occurring since 2005 (Bellwood et al., 2019). These frequent bleaching events reduced recovery intervals from bleaching episodes (Hughes et al., 2018). Coral bleaching events have recently shifted from average heat stress exposures of 4 to 12 °C-weeks to exceeding 24-35 °C-weeks in some reef locations (Hoegh-Guldberg 2011, Claar et al., 2018; Boyle et al., 2017; Brainard et al., 2018). The 2014-2017 global coral-bleaching event caused widespread mortality of corals and other reef-associated organisms over thousands of square kilometers (Eakin et al., 2019). Furthermore, bleaching events diminish coral recovery potentials such as reproduction, larval succession, and growth rates, which result in reduced species diversity, richness, and dynamics (Albright and Langdon, 2011; Albright and Mason, 2013; Anthony et al., 2008; Fordyce et al., 2019; Prada et al., 2017). Climate models have predicted future bleaching events with greater frequency and severity (more than 90% by 2030 and nearly all reefs by 2050) (Cai, 2014). To achieve better prediction, climate models require historical data on sea surface temperature, light intensity, salinity, nutrient level, species response, and other variables to identify patterns and relationships for coral bleaching (Eakin et al., 2018, Liu et al 2018). Therefore, geographical footprints of bleaching frequency, intensity, and their impact pattern at a broad geographical scale, particularly from the understudied locations, are crucial to predict future coral community structure against the increasing thermal stress.

### Accurate bleaching survey is important to capture differential species susceptibility patterns to bleaching (winners and losers)

Although global warming has triggered unprecedented mass coral mortality across the ocean; the extent of bleaching at different reef region varies due to differences in temperature magnitude, coral species composition, past bleaching history, hydrodynamics, and several other local environmental factors (Anthony et al., 2008; Hughes et al., 2018, 2003; Swain et al., 2016). In many instances, some corals tolerate or recover better from bleaching than others in every geographical region including India (Kumaraguru, 2003; Manikandan et al., 2014; Pisapia and Burn, 2016; Thinesh et al., 2019a, De et al. 2022, 2023). In India, studies are at a nascent state to identify the factors (symbiont types) responsible for differential bleaching patterns (for example Thinesh et al., 2019; Mote et al., 2021). Generally, massive taxa such as *Porites* spp. and some *Favidae* are more resistant to bleaching (Baird and Marshall, 2002). Massive coral taxa can live in the bleaching state for longer and recover better than their branching counterparts (Grottoli et al., 2014; Loya et al., 2001; Thinesh et al., 2019a). Historically the fast-growing branching taxa (*Acropora* and *Pocillopora)* are thermally sensitive and susceptible to heat stress-driven bleaching (Putnam et al., 2012). In contrast, slow-growing massive corals such as *Porites* spp. and some *Favidae* are considered better tolerant to bleaching (Baird and Marshall, 2002, Darling et al., 2012). During the bleaching events, massive coral taxa can live in the bleaching state for longer and recover better than their branching counterparts, but branching corals can only stay bleached for a short time before they die or recover (Grottoli et al., 2014; Loya et al., 2001; Thinesh et al., 2019a). This differential species-specific response is influenced by a combination of factors, including microbiomes, types of symbionts, coral life-history traits, tissue thickness, investment in energy storage, colony size, growth rate, protein content in the mucus, coral species composition, past bleaching history, hydrodynamics (Berkelmans and van Oppen, 2006; Loya et al., 2001; McClanahan et al., 2004, Anthony et al., 2008; Swain et al., 2016). These factors significantly influence their susceptibility to bleach mortality and recovery in individual reefs, determining their future community structure (Camp et al., 2018; Hughes et al., 2003). Therefore, data on coral community composition, taxon-specific susceptibility, and resilience patterns to bleaching events are essential for every reef location (Cornwell et al., 2021; Edmunds, 1994; Thinesh et al., 2019a, Dixon et al., 2022). Coral bleaching survey is a widely used approach to record this pattern in the field. Therefore, surveying efforts must be uniform to use the data for local and global scale comparison. During short-term bleaching events (DHWs > 4), one-time surveys could document the complete bleaching impact as bleaching windows narrowed. In cases of long-term bleaching events, repeated surveys are necessary to document the impact of heat stress accurately, especially to determine the winners and losers (Heron et al., 2016, Danielle et al., 2018). If repeated surveys are not feasible, a study must include DHWs for their locations, the bleaching proportion of each coral genera, which allow us to validate and compare bleaching effects across studies and regions (Claar et al., 2018).

In India, the first mass coral bleaching episode was reported in 1998. Since then, many surveys have documented the bleaching episodes, impact, and species susceptibility pattern from multiple bleaching events across different reef sites. As major reefs are separated geographically, most available bleaching data across this 23-year timespan are patchy, limited to one-time surveys, or restricted to a single bleaching episode from a specific location. The scattered data across reef sites hinder the understanding of species susceptibility patterns and bleaching impact. In this study, we examined the available published research on the thermal history, bleaching intensity, mortality, and species susceptibility pattern from 1998 to date; (i) to create a comprehensive database of bleaching episodes in Indian reefs; (ii) to assess bleaching percentage in respective to DHWs; iii) to identify susceptible and resistant coral genera during long-term (DHWs > 4) and short-term bleaching events DHWs < 4; (iv) to highlight gaps in monitoring efforts; and iv) to provide uniformed future survey recommendations.

## Methods

Preferred Reporting Items for Systematic Reviews and Meta-Analyses (PRISMA) was adopted for the current review, using different scholarly databases, namely Scopus™, Web of Science™, and Google Scholar™, as these databases are robust and cover ecology, environmental studies, and oceanography. PRISMA is a published standard to conduct a systematic literature review, also suitable for the environmental management field because it clearly defines the research questions for a systematic review (Moher et al., 2009; Sierra-Correa and Cantera Kintz, 2015). We also carried out manual searches on several other journal databases such as Science Direct, Taylor Francis, Springer, and Sage, considering that they contain peer-reviewed journal articles related to the field of coral reefs. Primary literature (concerning Indian coral reefs and coral bleaching) published during the timeline, 1998 to 2021 was extracted using the following keywords: (coral reef OR tropical reef OR coral OR Scleractinia OR hard coral OR stony corals OR octocorals OR soft corals OR Coral bleaching OR Coral mortality Or climate change OR global warming OR temperature rise OR SST anomaly OR el-Nino OR la-Nina OR Indian reef OR Gulf of Kachchh OR Malvan Marine Sanctuary OR Grande Island OR Angriya bank Or Netrani Island OR Lakshadweep OR Palk Bay OR Gulf of Mannar OR Andaman Or Nicobar Or Bay of Bengal Or Arabian Sea. Only peer-reviewed articles were considered for the present review as the primary sources that offer empirical data to maintain both consistency and quality. A careful cross-referencing was also conducted to include any articles missed during the literature search. It should be noted that the review only focused on articles published in English. Other than that, only studies conducted in the Indian territorial water were selected because they are in line with the review’s objective. Our search retrieved 210 articles from the databases at the first stage of the systematic review process. A thorough screening was conducted to remove duplicate reports. Based on the degree of relevance, individual papers were then eliminated by ‘title’ or ‘abstract’ alone, or by accessing the entire article. A total of 36 published research papers were reviewed as part of this effort. We avoided reports published in the newspapers as it has no data on species susceptibility, percentage of bleaching, and mortality except the evidence of bleaching. Fig. 1 represents the flow diagram for the PRISMA analysis.

**Fig. 1.**
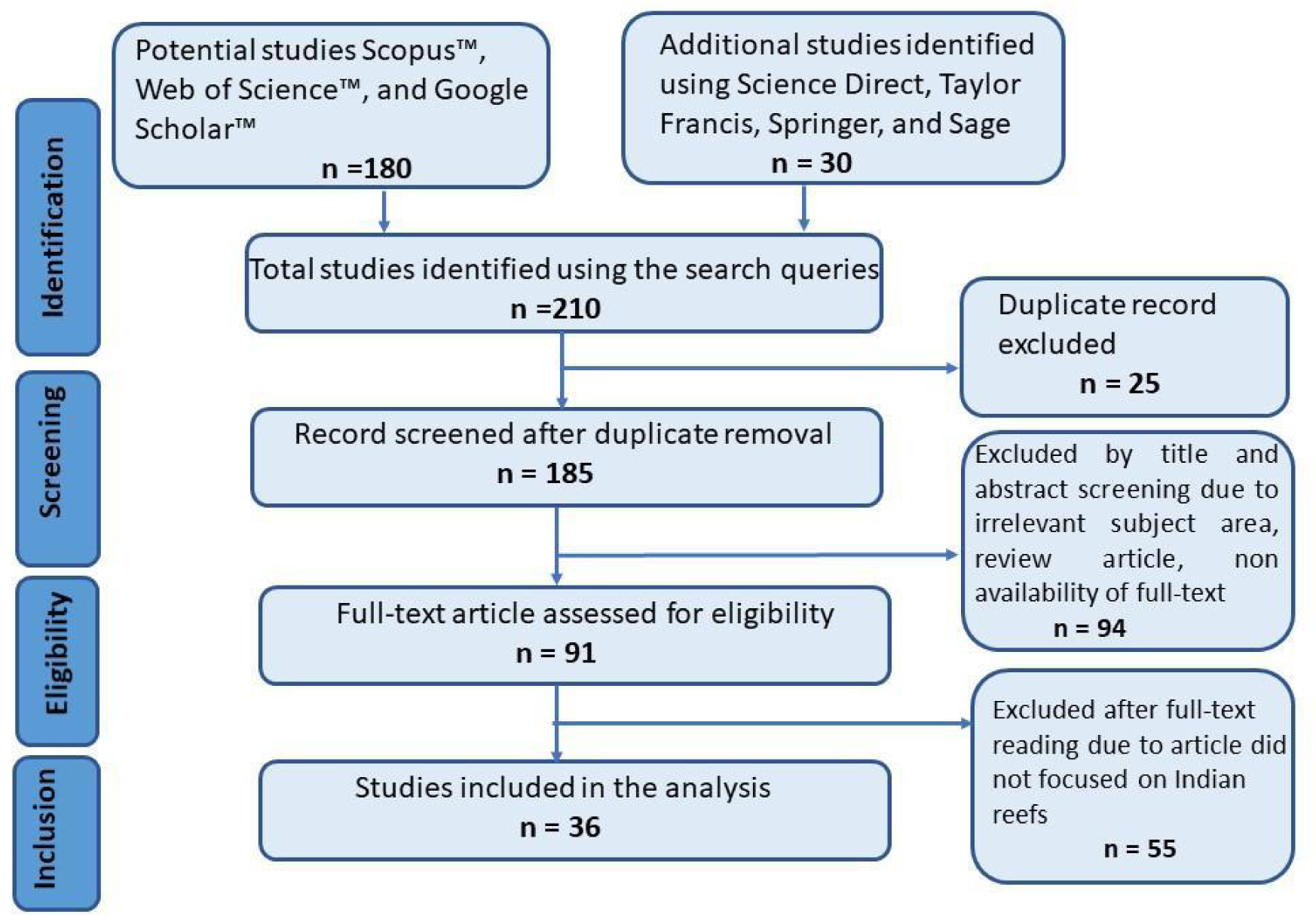
Flow diagram of the PIRSMA analysis (adapted from Moher et al., 2009)

## Results

**Coral reef distribution in India and the bleaching reports:**

India has four major coral reef regions and a few patchy reefs. Major coral reefs are located in the i) Gulf of Kachchh, ii) Gulf of Mannar biodiversity reserve, iii) Lakshadweep Islands, and iv) Andaman and Nicobar Islands separated geographically. In addition, there are three patch reefs 1) Palk Bay, 2) Malvan Marine Sanctuary, 3) Grande Islands, and a few patches at different locations as follows Ratnagiri, Redi, Netrani, Quilon in the Kerala coast to Enayem in Tamilnadu, Parangipettai (Porto Novo), south of Cuddalore, and Pondicherry (De et al., 2017). The estimated coral reef area in India is around 2,375 km^2^ (DOD and SAC 1997). These coral reefs harbor a total of 585 species belonging to 108 genera and 23 families (De et al., 2020). Studies have documented a steady rise in SST anomaly and thermal stress in Indian reefs (Arora et al., 2019, 2021b, De et al., 2022). The spatial distribution of the warmest SST, SST anomaly, and DHWs data for each reef environment indicates that corals experienced thermal stress and likely mass bleaching (DHWs ≥ 4) (Arora et al., 2019).

### A history of bleaching events and their mortality

Indian reefs experienced three long-term (major) and ten short-term (Minor) bleaching episodes within the last 23 years. All three major bleaching events (1998, 2010, and 2016) affected all four major reefs, which caused mortality. Ten short-term bleaching events are scattered among major reefs and patchy reefs out of 13 recorded bleaching events from seven reef sites. The Gulf of Mannar experienced many bleaching events compared to other reef sites (Fig 3). The bleaching percentage and mortality report on bleaching events is given in Table 1.

**Fig. 2.**
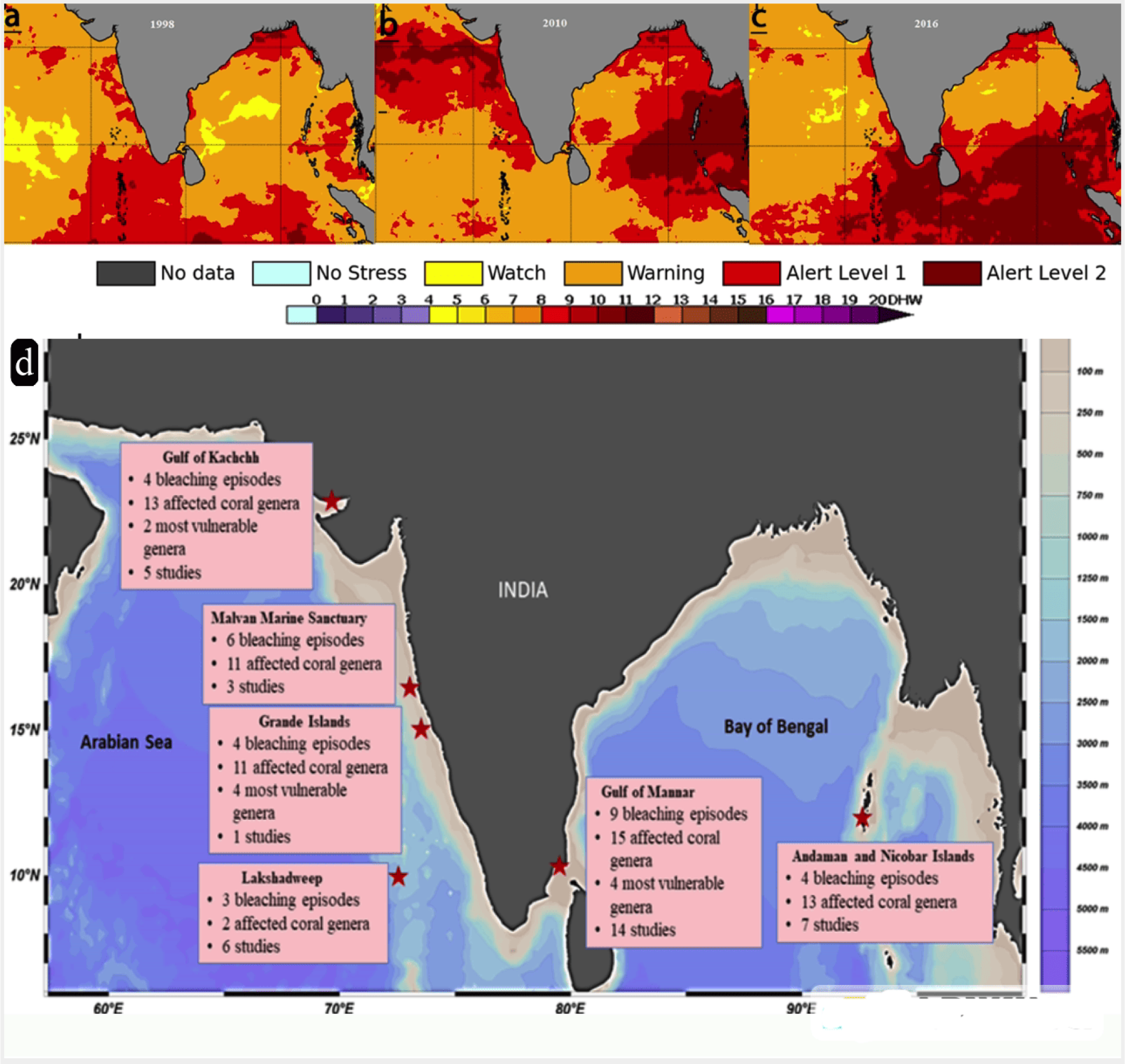
Bleaching area alert annual maximum a) 1998; b) 2010; c) 2016 (Data and Plot: NOAA coral reef watch V3.1). d) Geographical distribution of the major Indian coral reefs with reported coral bleaching episodes

**Fig. 3.**
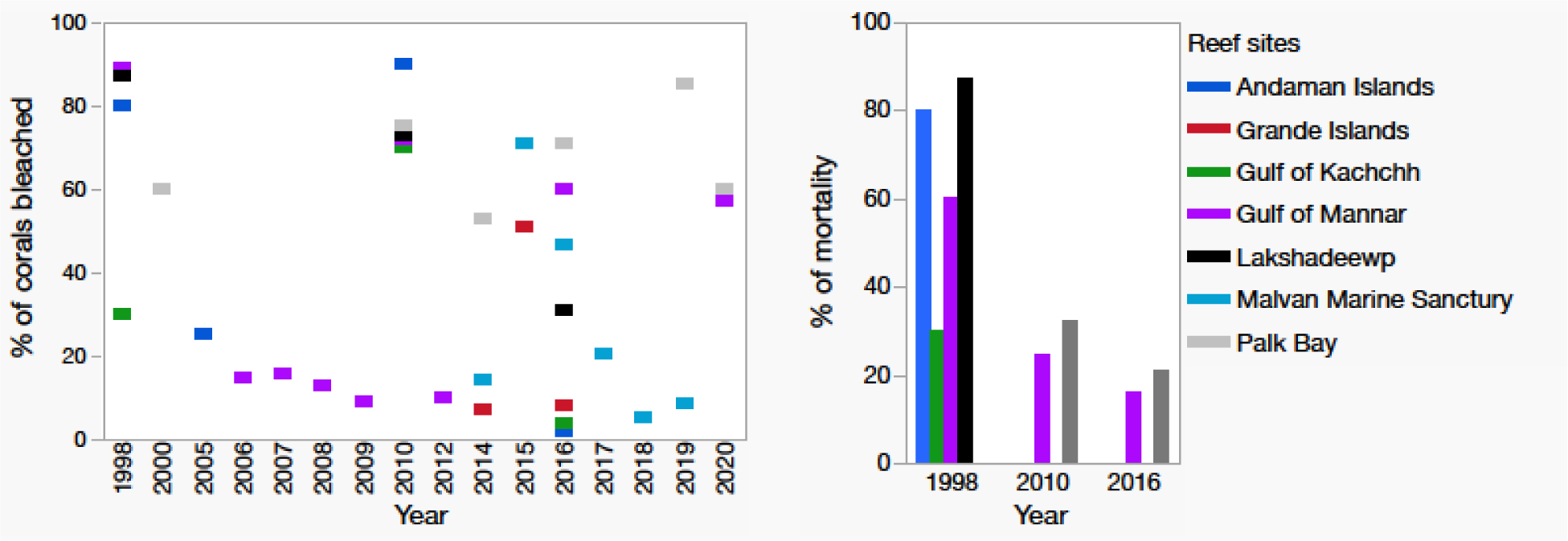
(A) Several recorded bleaching events and their percentage from the major and patchy reefs. (B) Observed bleaching-induced mortality events in seven Indian reefs (we used the highest mortality percentage report for representation. Details are given in the Table 1.)

**Table 1:**
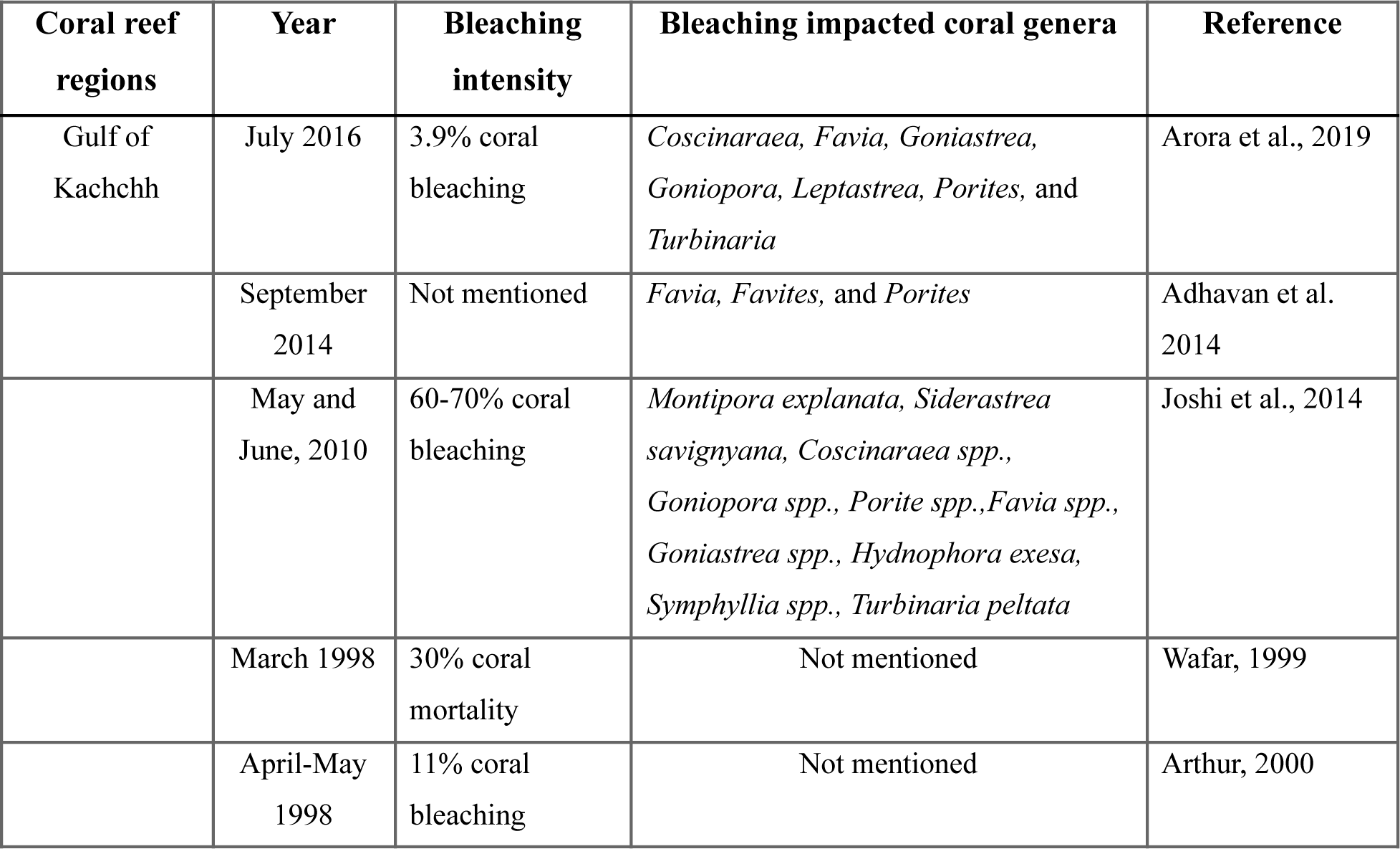
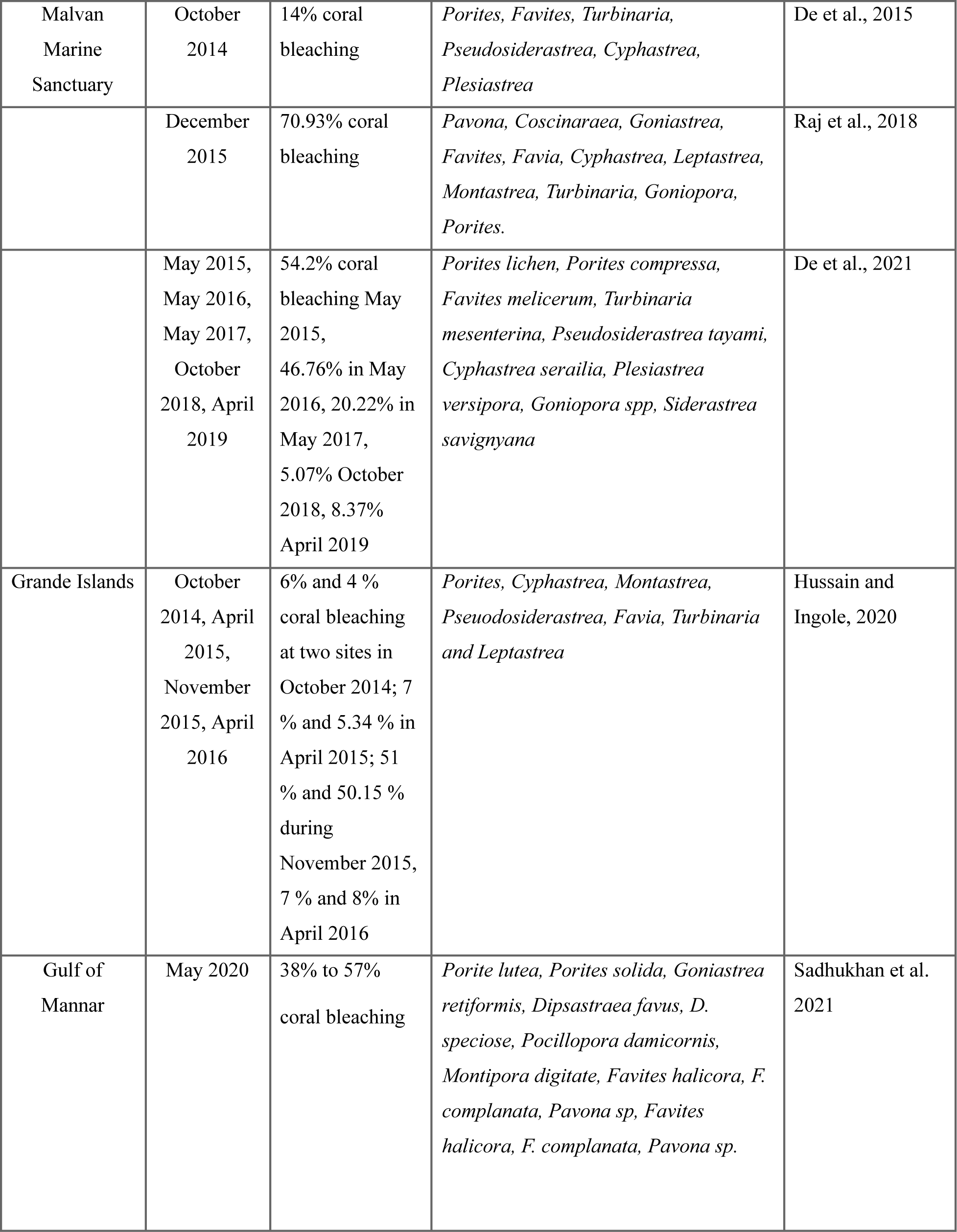
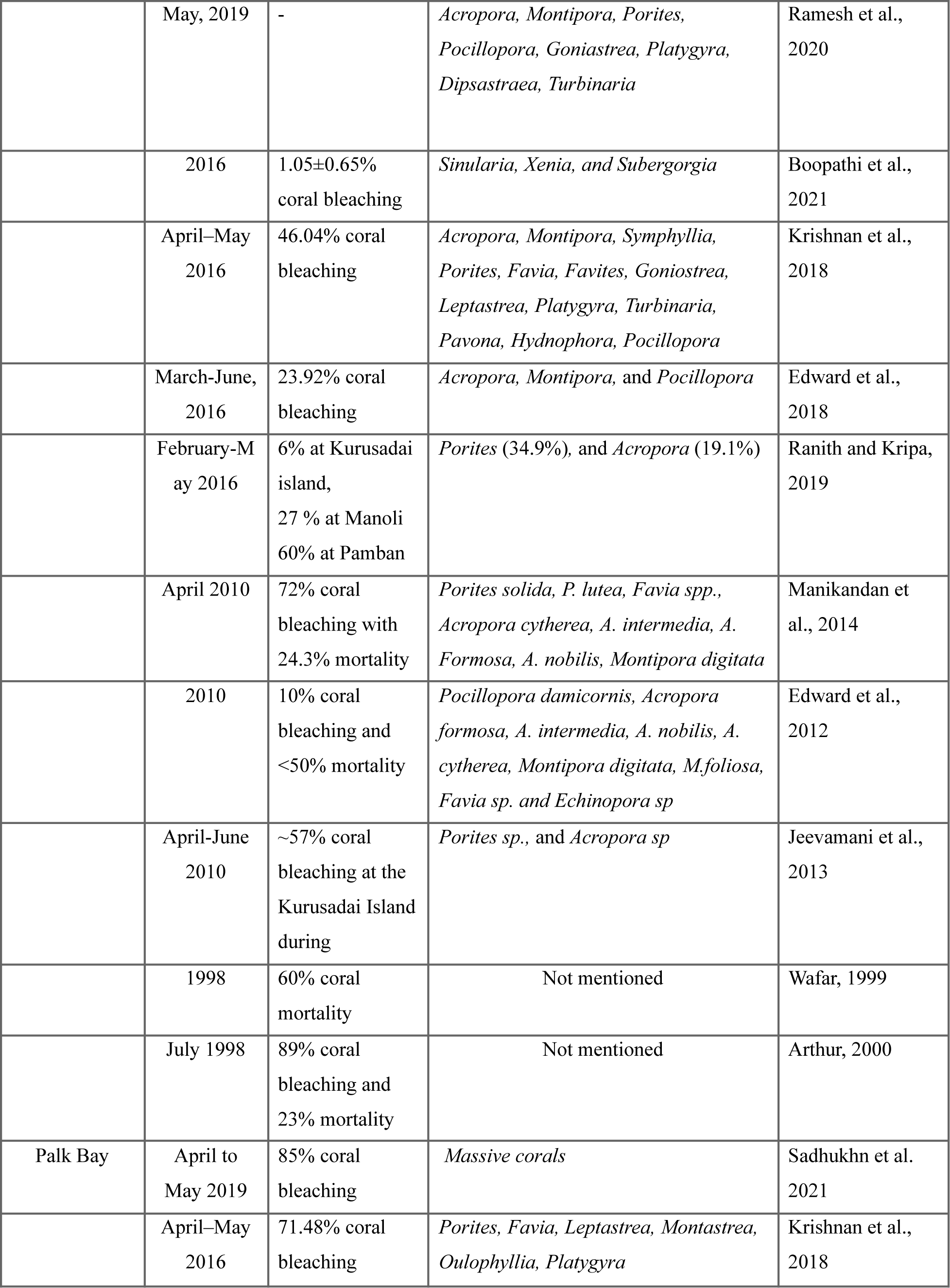
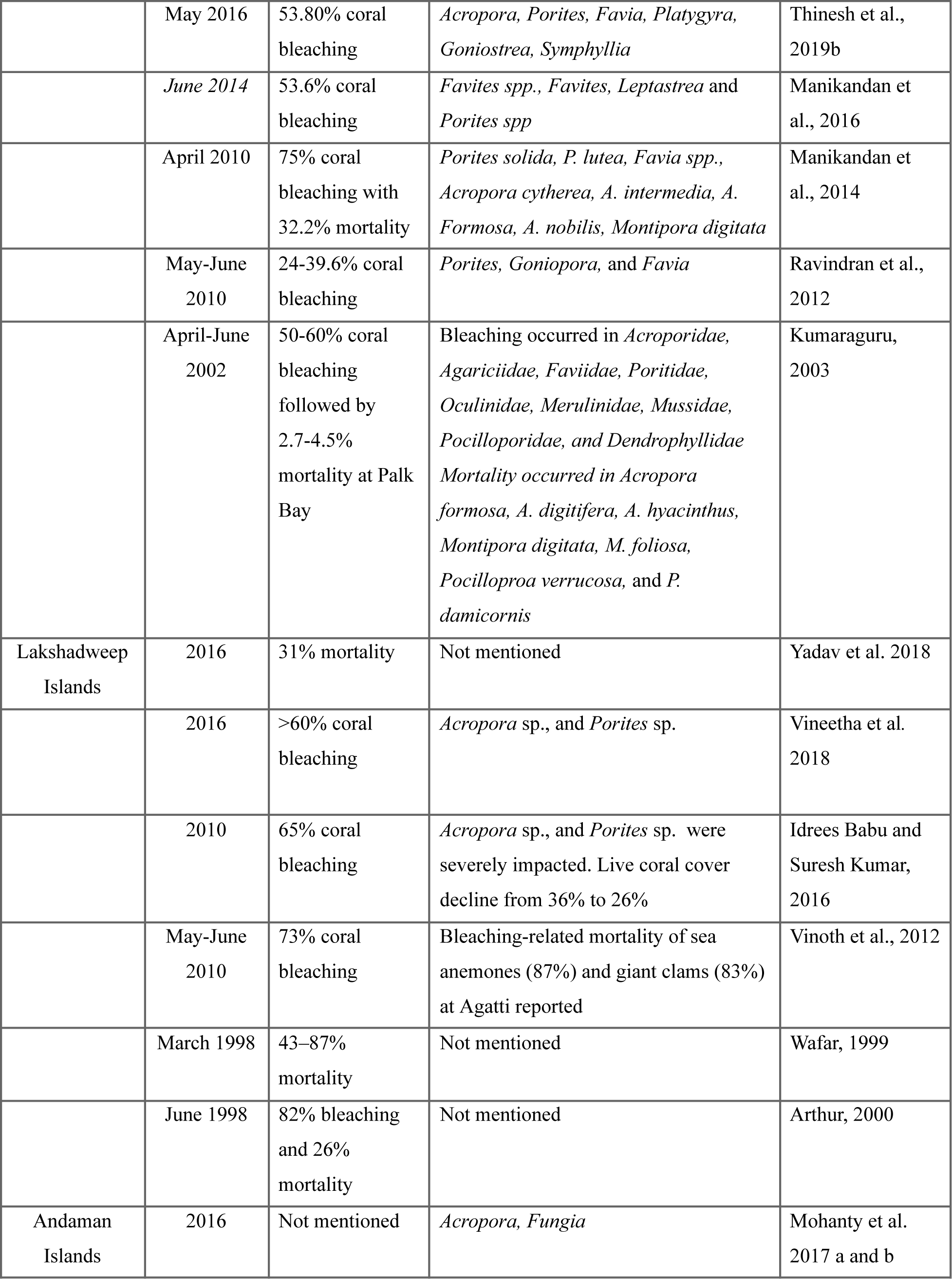
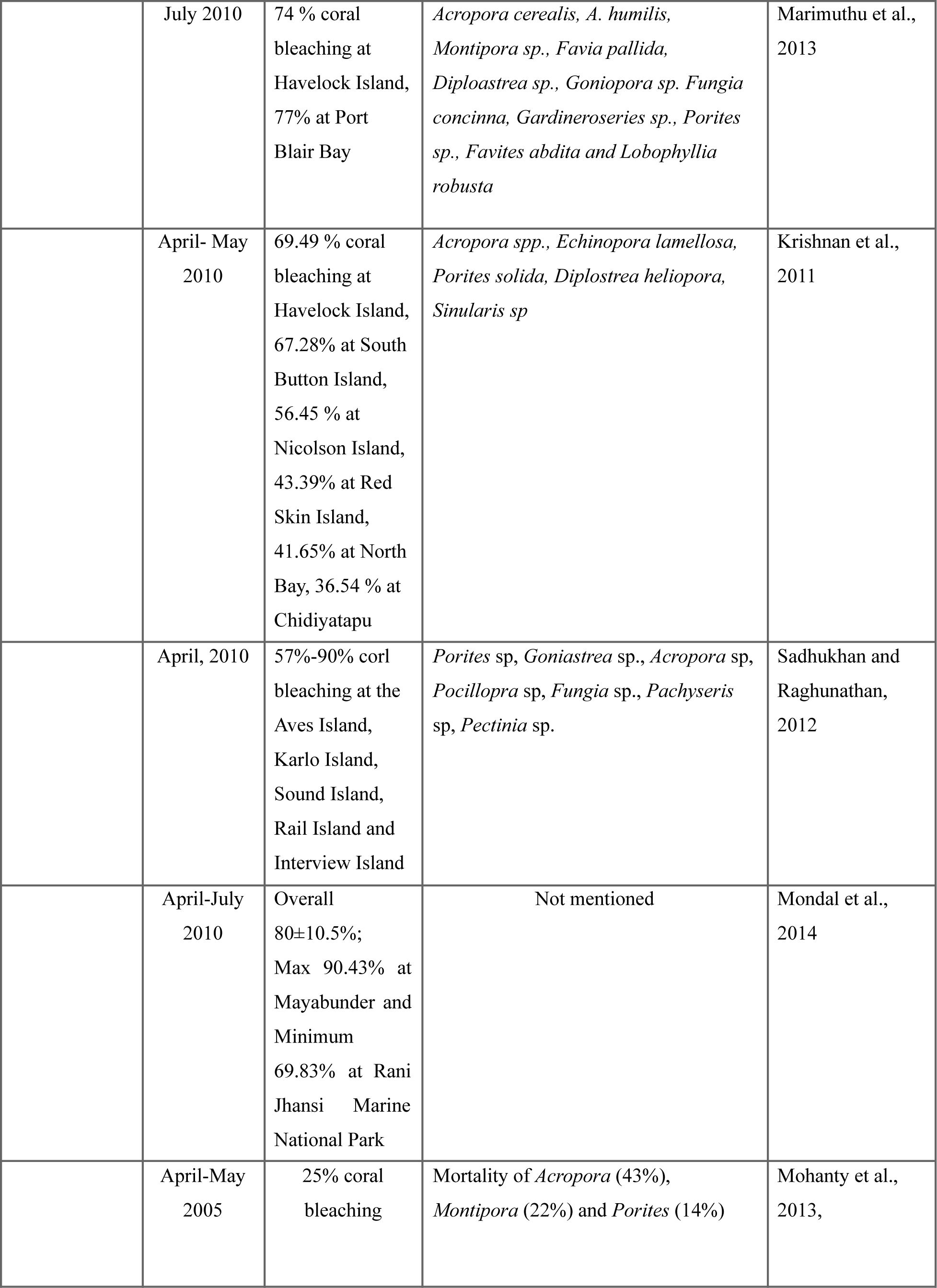
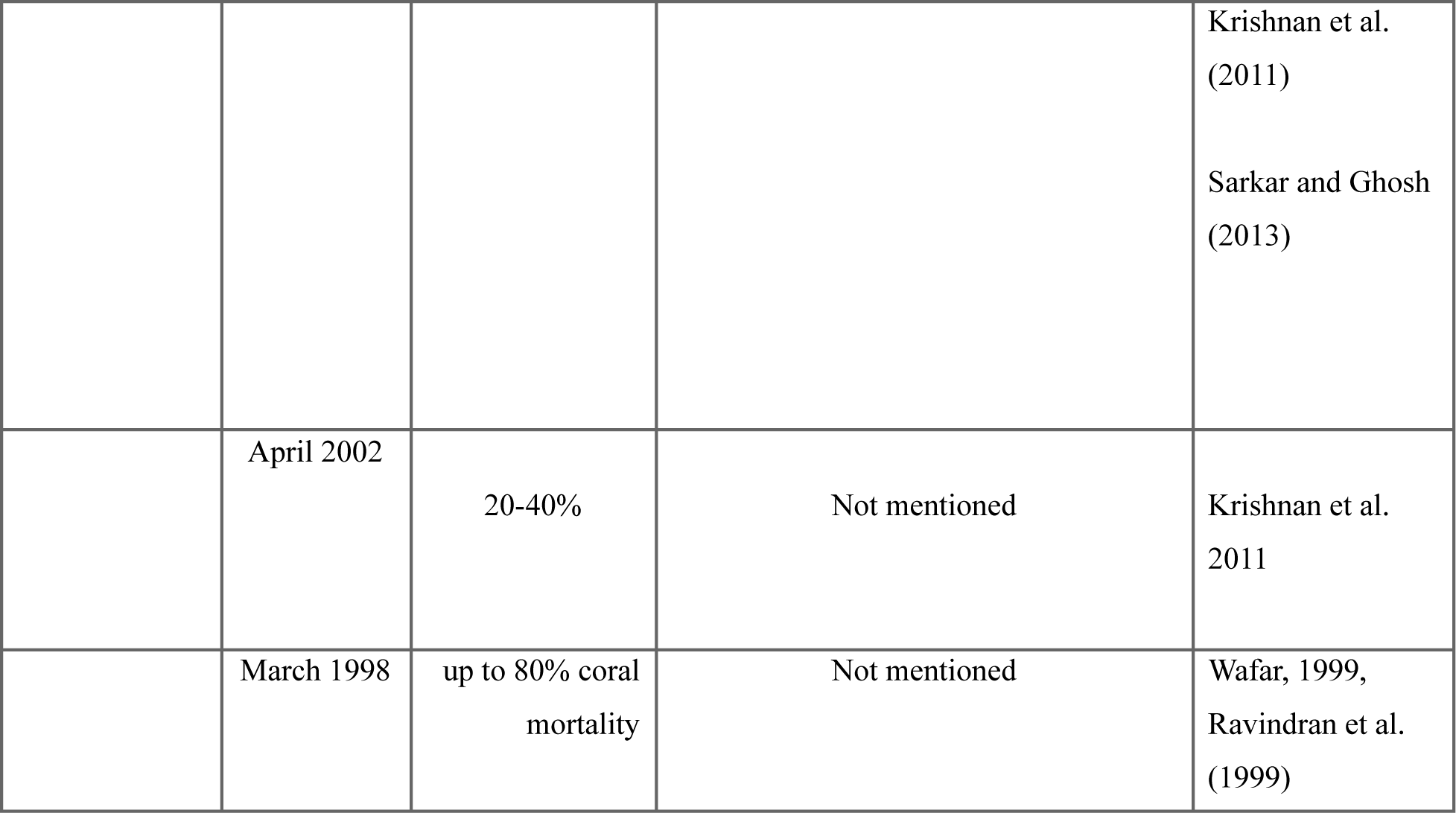
Occurrence of different coral bleaching episodes in major coral reefs in Indian territorial water.

### Experienced degree heating weeks and reported bleaching percentage from four major reefs

Indian coral reefs were exposed to different degree heating weeks (DHWs; ^°^C-weeks)-an indices for assessing thermal stress based on satellite-derived SST data, and their bleaching percentage differ among sites for all three major bleaching events. For 1998, 90% of reefs experienced bleaching in the Gulf of Mannar, with 23% mortality to DHWs 6.7. In Andaman, 80% of reefs experienced bleaching to DHWs 4.9. In Lakshadweep, 82% of reefs experienced bleaching with 26% mortality to DHWs 6.17, while in the Gulf of Kachchh, 11 to 30% of reefs experienced bleaching to DHWs 2.16. For 2010, In Andaman and Nicobar, 36.5% to 77% of reefs experienced bleaching to DHWs 11.7. Whereas, 10% to 72% of reefs experienced bleaching, and 24% mortality to DHWs 5.6 in the Gulf of Mannar. In the Gulf of Kachchh, 60-70% of reefs experienced bleaching to 11 DHWs. In 2016, In Andaman & Nicobar reef experienced DHWs 7.21 to 9.5, but this site did not report the bleaching. In the Gulf of Mannar, 23% to 60% of reefs experienced bleaching to DHWs 7.6 (41 days), while in the Gulf of Kachchh, 3.9% of reefs experienced bleaching to DHWs 5.57. Above mentioned information is graphed in Fig 4, and the details of thermal stress events on Indian reefs are given in the Table 2 and 3. Daily trends of mean SST and SST anomaly for the period 2010 to 2021 in major Indian coral region (Data source: NOAA CRW v 3.1) were represented in the Supplementary Fig S1-S5.

**Fig. 4.**
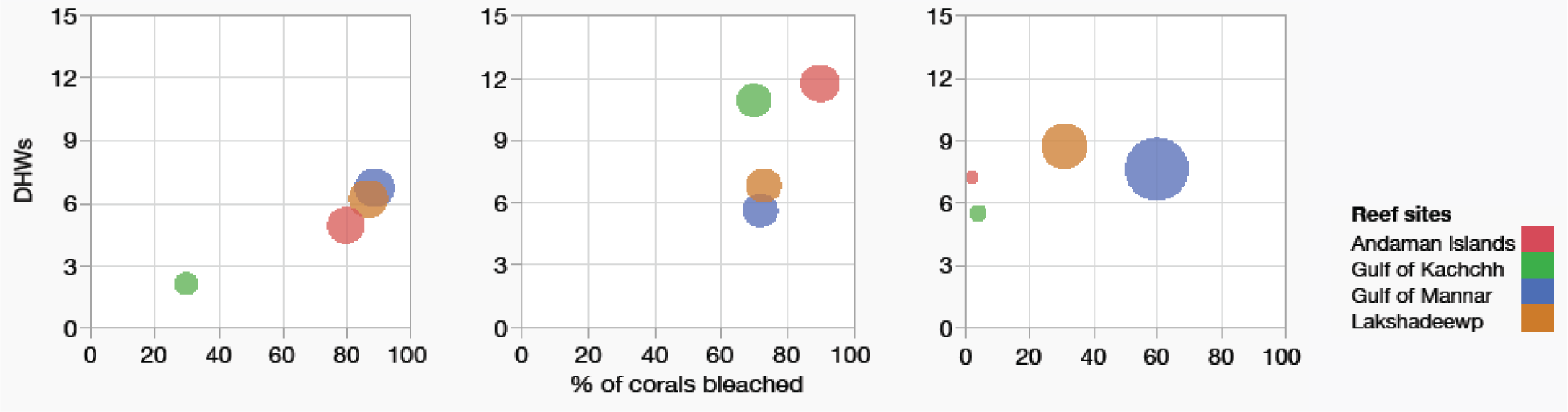
Degree heating weeks and recorded bleaching percentage to three major bleaching events from major Indian coral reefs

**Table 2:** Warmest month’s sea surface temperature (SST), warmest month, SST anomaly, duration of thermal stress and degree heating weeks for mass coral bleaching years in the Indian coral reef regions based on Arora et al. 2019.

| Year | Coral bleaching indices | Andaman | Nicobar | Lakshadweep | Gulf of Mannar | Gulf of Kachchh |
| --- | --- | --- | --- | --- | --- | --- |
| 1998 | Warmest month SST (°C) | 30.66 | 30.69 | 30.71 | 30.84 | 29.58 |
|  | Warmest month SST anomaly (°C) | 0.76 | 0.87 | 0.81 | 0.61 | 0.23 |
|  | Duration of thermal stress (days) | 61 | 67 | 68 | 67 | 37 |
|  | Total no. of continuous days (%) | 39.87 | 43.79 | 44.44 | 43.79 | 24.18 |
|  | Degree heating week (°C) | 4.91 | 6.43 | 6.17 | 6.77 | 2.16 |
| 2010 | Warmest month SST (°C) | 31.43 | 31.26 | 30.77 | 30.87 | 30.44 |
|  | Warmest month SST anomaly (°C) | 1.53 | 1.44 | 0.87 | 0.64 | 1.09 |
|  | Duration of thermal stress (days) | 87 | 85 | 73 | 48 | 91 |
|  | Total no. of continuous days (%) | 56.86 | 55.55 | 47.71 | 31.37 | 59.48 |
|  | Degree heating week (°C) | 11.72 | 12.24 | 6.8 | 5.06 | 10.98 |
| 2016 | Warmest month SST (°C) | 31.03 | 31.2 | 30.86 | 31.21 | 29.91 |
| t | Warmest month SST anomaly (°C) | 1.13 | 1.38 | 0.96 | 0.98 | 0.56 |
|  | Duration of thermal stress (days) | 53 | 67 | 91 | 63 | 70 |
|  | Total no. of continuous days (%) | 34.42 | 43.51 | 59.09 | 40.91 | 45.45 |
|  | Degree heating week (°C) | 7.21 | 9.51 | 8.71 | 7.69 | 5.57 |

**Table 3:** Warmest months, thermal thresholds, and bleaching thresholds of Indian coral reefs regions by using NOAA OISST data from 1982 to 2018 based on Arora et al. 2019; and De et al. 2021.

| Coral reef regions | Warmest month | Warmest quarter | Thermal threshold (°C) | Bleaching threshold (°C) |
| --- | --- | --- | --- | --- |
| Gulf of Kachchh | June | March-July | 29.35 ± 0.45 | 29.85 |
| Malvan | May | March-May | 29.39° C + 0.49 | 29.89 |
| Lakshadweep | May | March-May | 29.90 ± 0.49 | 30.4 |
| Gulf of Mannar | April | March-May | 30.23 ± 0.39 | 30.73 |
| Andaman | May | April-June | 29.90 ± 0.58 | 30.4 |
| Nicobar | May | March-May | 29.83 ± 0.63 | 30.33 |

### Species susceptibility responses - winners and losers

To rank the species’ susceptibility to bleaching events, we have selected two major bleaching events (2010 and 2016) and one short-term bleaching event from three major reef regions (Gulf of Mannar, Gulf of Kachchh, and Andaman reef). These reefs had species response details to 2010 and 2016 major bleaching episodes and one short-term bleaching episode, as the 1998 bleaching event does not have species response details. Four coral genera in the Gulf of Mannar, *Acropora, Pocillopora, Montipora,* and *Porites* were bleached to short-term bleaching events (lowest DHWs - 2). In addition, *Echinopora,* and *Favia* were bleached in 2010 (DHWs, 5.6). In 2016, When DHWs increased to 7.69, susceptibility differences diminished among species. In the Gulf of Kachchh, *Porites,* and *Favia* were bleached to short-term 2010 bleaching. In addition, *Goniopora, Goniastraea, Leptastrea,* and *Gascinaraea* were bleached in 2016 (DHWs-5.57, WM 29.9). The disparity disappeared among coral genera to 2010 long-term bleaching episode (DHWs-10.98, WM-30.44). In Andaman reefs, *Acropora, Montipora,* and *Porites* were bleached to all three bleaching events. *Fungia* was bleached during 2016 in addition to short-term events. Again, the disparity in susceptibility disappeared when DHWs extended to above ten weeks (DHWs 11-12). Bleached coral genera details against DHWs from different reef site details are given in Fig. 5.

**Fig. 5.**
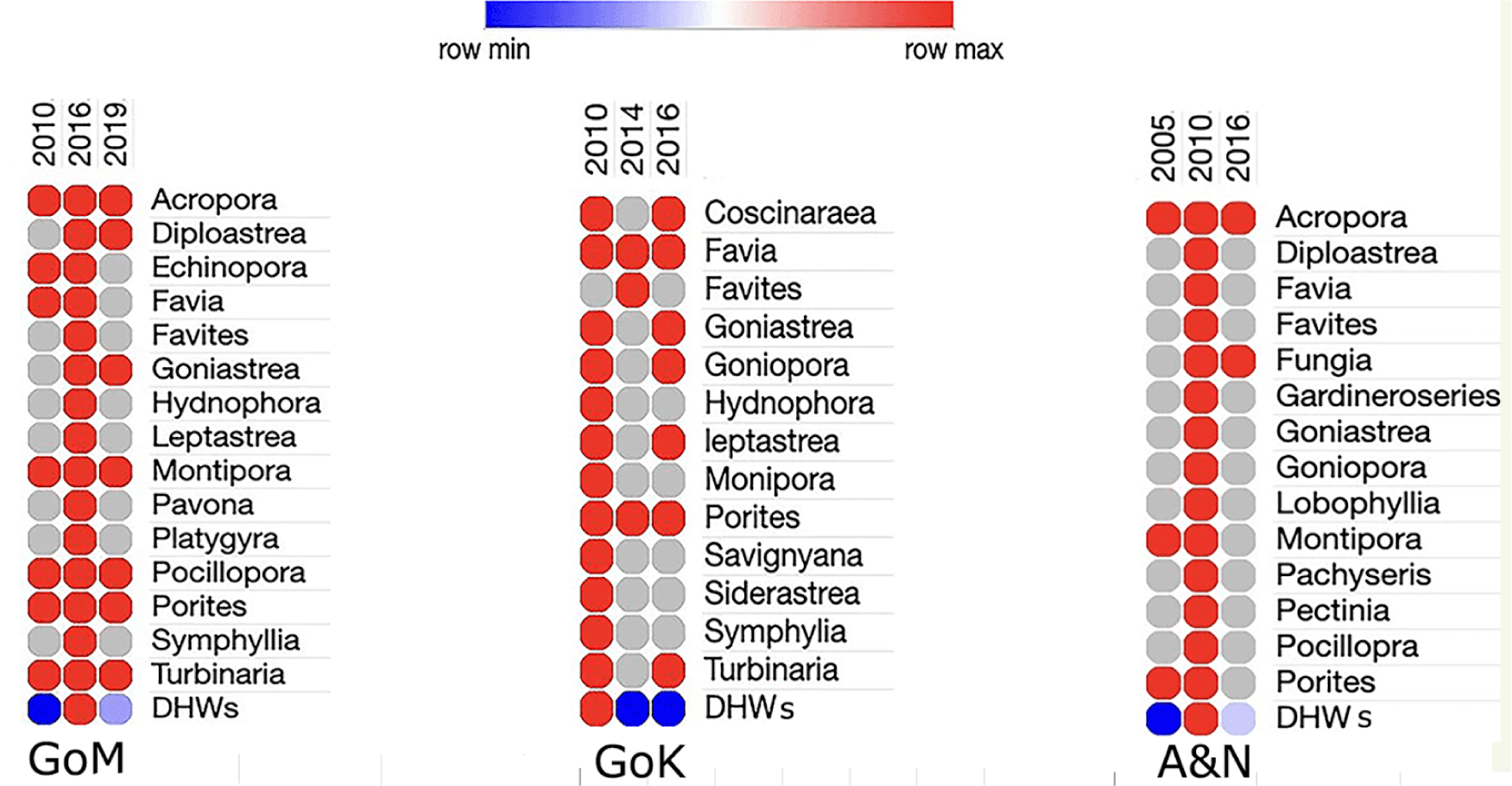
Bleaching response observed in coral genera to one short-term and two long-term (Major) bleaching events from left to right 1) Gulf of Mannar (GM), 2) Gulf of Kachchh (GoK), and 3) Andaman and Nicobar Islands (A&N).

### Contrasting conclusions on species susceptibility to the 2016 long-term bleaching event in the Gulf of Mannar and Palk Bay reef

In the Gulf of Mannar, three studies documented the impact and species susceptibility pattern during the 2016 (DHW= 7.6, highest temperature 31.2°C) long-term bleaching event. All these three surveys are independent one-time surveys. These three studies created excellent data, but their conclusions differ on the bleaching percentage and species susceptibility ranking. Krishnan et al. (2018) found that 46% of reefs experiencing bleaching and 68% of bleached corals were massive forms from the Tuticorin group of islands. They found that 22 out of the 26 massive forms were bleached, but the *Acropora* corymbose (ACC), digitate (ACD), and encrusting coral (CE) forms had not bleached in any of their study sites. Based on the findings, the study concluded that branching corals might have adapted to bleaching as they have experienced repeated bleaching events in the Gulf of Mannar.

In contrast, Edward et al. (2018) found the opposite pattern. They reported that 23.92% of reefs experienced a 2016 bleaching episode (21% of reefs in the Tuticorin reef). They found higher susceptibility and mortality in branching corals than in the massive forms (mortality and bleaching percentage not provided) (Edward et al., 2018). This study concluded that fast-growing branching corals (fast-growing coral forms, *Acropora*, *Montipora*, and *Pocillopora* are more susceptible to bleaching than the boulders (*Porites, Favia,* and *Favites*). Another study reported more susceptibility in massive forms than in branching forms (Ranith and Kripa, 2019). In the case of Palk Bay, two studies contrast one another. Krishnan et al. (2018) reported that 71% of reefs experienced bleaching, and the highest susceptibility was in *Porites*, *Favia, Leptastrea, Montastrea, Oulophyllia,* and *Platygyra*, wherein the highest vulnerability was observed in *Acropora* than the massive life forms by different study (Thinesh et al., 2019b).

**Fig. 6.**
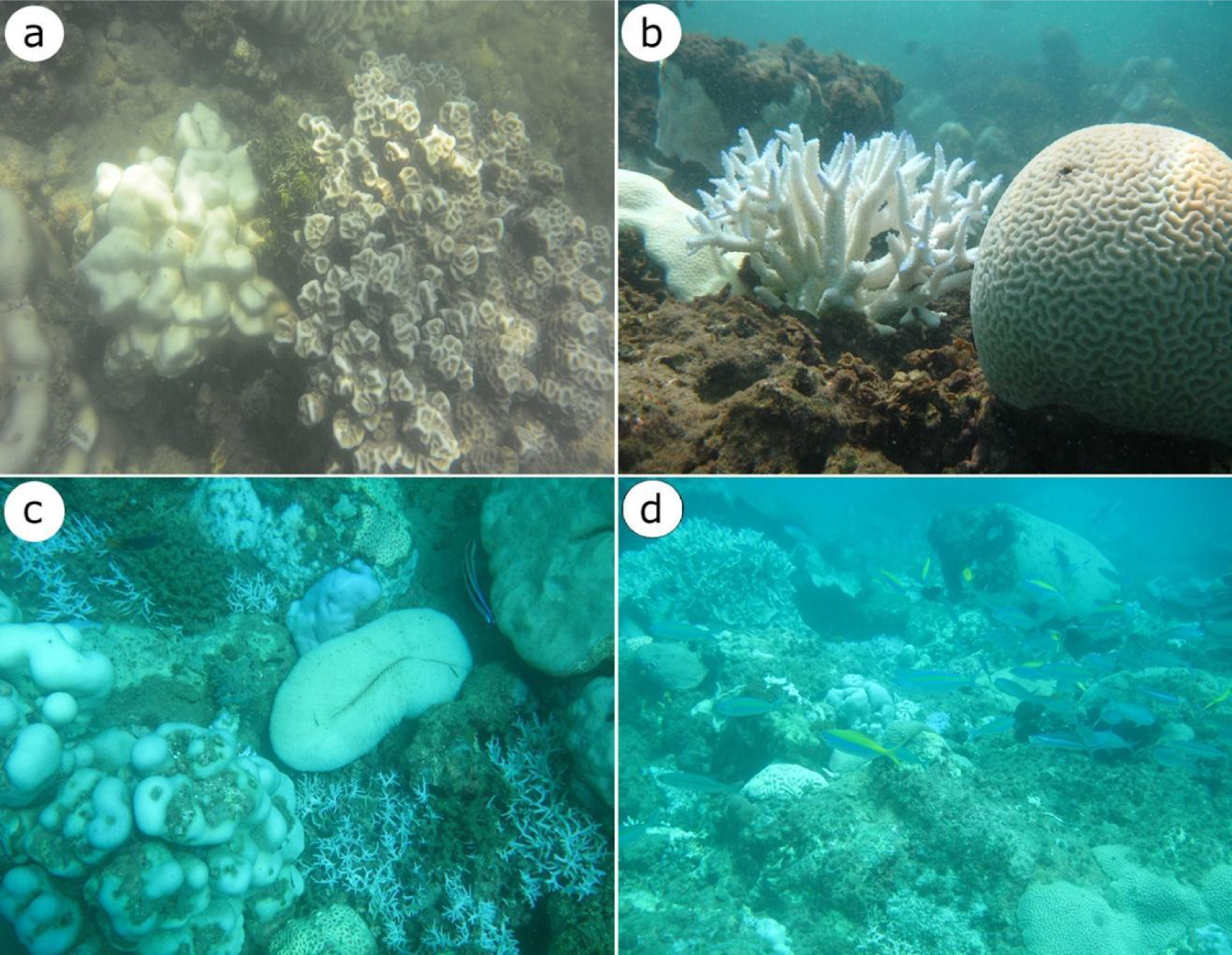
Differential coral bleaching patterns observed from Indian coral reefs. A) Partially bleached *Porites* near to healthy *Favites b)* Bleached *Acropora*, *Favites*, partially bleached *platygyra* and bleached dead *Porites* partially covered by algae c) Bleached dead *Acropora* and bleached *Fungia*, *Goniastrea* and *Porites* d) Bleached dead Acropora along with bleached other coral genera.

## Discussion

Global data shows that the degree of bleaching differs at different geographical locations. Similarly, based on the literature survey, we found variation in bleaching percentage and species response among Indian reefs. The observed bleaching intensity variation between study locations is likely due to the coral composition type, type of symbionts, depth, time of exposure, and various other factors (Hoogenboom et al., 2017). The Gulf of Kachchh is least susceptible to major bleaching events among all major reefs. For instance, in 2016, DHWs in the Gulf of Kachchh reached 5.57 weeks; only 3% of reefs experienced bleaching, while in the case of the Gulf of Mannar during 2010 (DHWs-5.6, 2010), it was 10 to 70%, and in Andaman, it was ∼ 80 (DHWs-4.91, 1998). This differential susceptibility between locations is likely due to differences in the taxonomic composition. (Marshall and Baird, 2000). The highest proportion of massive forms and the absence of sensitive coral genera (*Acropora, Pocillopora*) in the Gulf of Kachchh likely favors the better resilient to bleaching as massive forms are stress-tolerant (Darling et al., 2012, Yohesh et al, 2014). The physiological mechanism that leads to higher resistance in massive corals is unknown to Indian reefs. However, attempted studies in other locations found the highest metabolic rate (Gates and Edmunds, 1999), tissue thickness (Loya et al., 2001), mass transfer rates (Nakamura and Van Woesik, 2001), light absorbing capacities (Fabricius, 2006), heterotrophy (Grottoli et al., 2006), and symbiont association (Baker et al., 2004) favors a role in resistance.

Most bleaching studies have found that branching coral genera such as *Acropora* and *Pocillopora* are more susceptible to bleaching than slow-growing massive corals (Baker et al., 2004; Loya et al., 2001; Marshall and Baird, 2000; McClanahan et al., 2004), except few studies reported contrasting patterns (Guest et al., 2012; Krishnan et al., 2018). This observation has led to the widespread classification of fast-growing branching species as “losers” in the face of global warming (Loya et al., 2001), at least in the short term (Van Woesik et al., 2011). Likewise, our data shows the highest susceptibility in *Acropora* at every location except in the Gulf of Kachchh, where the *Acropora* population diminished for unknown reasons. In contrast, *Porite’s* susceptibility to bleaching is also invariably recorded in short and long-term events across the site. We infer that *Porites* and massive colony types exhibit earlier partial bleaching (pale coloration). However, they remain in the bleaching state for the entire bleaching event and recover, unlike their sensitive counterparts who underwent mortality. Since existing data undifferentiated partial/complete bleaching, the *Porites* fall under susceptible genera. This pattern suggests that if the survey happens in the first week, one could expect partial bleaching in the *Porites* colonies even before the susceptible branching forms start to pale. In the Gulf of Mannar, *Montipora,* and *Pocillopora* are the other three genera equally susceptible to short-term events. Other coral genera that have only bleached during long-term bleaching events are given in the graph. Similar to bleaching susceptibility, coral taxa differ in their recovery pattern (Baird & Marshall, 2002). A study from the Great Barrier Reef (GBR) recorded the highest recovery in massive corals than the branching corals. For instance, a tagged colony study found that 88% of *Acropora hyacinthus,* and 32% of *A. millepora* colonies died compared to 13% of *Platygyra daedalea* and 0% of *Porites lobata* (Baird & Marshall, 2002). Although few studies looked at the recovery pattern in coral genera immediately following bleaching, the study found the highest recovery in the massive type than the branching corals (Manikandan et al., 2016b; Thinesh et al., 2019b). In support of that, a large-scale survey in the Gulf of Mannar after the mass bleaching event documented an increased massive coral cover in the coral community composition as branching forms were presumed to have died as a result of bleaching. (Raj et al., 2021). In India, studies are beginning to understand factors leading to the highest resistance among specific coral taxa and growth forms (Thinesh et al., 2019). We assume this susceptibility pattern in coral genera may be associated with differences in the type of symbiont hosted (Berkelmans and van Oppen, 2006), pre-bleaching history, metabolic rates (Gates and Edmunds, 1999), tissue thickness (Loya et al., 2001), and heterotrophic feeding capacity (Grottoli et al., 2006).

We found a susceptibility pattern among coral genera to short-term bleaching events. However, the disparity diminished to long-term bleaching events similar to the study, which reported no difference in species susceptibility to prolonged bleaching events in the GBR (Hughes et al., 2018). In addition, the scientific community hypothesized that recurrent bleaching episodes favor coral to adapt or acclimatize to future bleaching episodes. Studies have indicated that previous recurrent bleaching exposure protects corals from subsequent episodes (Carilli et al., 2012; Guest et al., 2016; Palumbi et al., 2014; Trapon et al., 2011). In contrast, one large-scale study from the GBR found no evidence of acclimation or adaptation in the corals exposed to earlier bleaching events (Hughes et al., 2018). Our analysis of available literature shows no evidence of protective effects from previous bleaching history in Indian coral reefs. For instance, the Gulf of Mannar reefs experienced eight bleaching episodes, including three long-term and five short-term. However, 23 to 60% of reefs experienced bleaching in 2016 (DHWs-7.69), and the same coral genera show the highest susceptibility (Raj et al.2021).

Our analysis of species susceptibility reports for 2016 long-term bleaching events from the Gulf of Mannar and Palk Bay revealed contrasting conclusions on species susceptibility ranking. Then, we examined why the study found a contrasting pattern of results even though they surveyed the exact locations. This pattern may be attributed to the colonies they encountered being different from one another, which might have experienced different thermal histories. However, we assume that the survey timing is most likely to influence the pattern difference between the studies similar to the bleaching surveys undertaken during the 1997/1998 El Nino (surveyed 10, 14, 20, 28, and 40 weeks after bleaching began; Marshall and Baird, 2000) as well as the 2015/2016 El Nino from Kiritimati Island (Claar and Baum, 2019). In GBR Marine Park, a survey during the 6th week found the highest susceptibility to the *Acropora hyacinthus* population, while the massive reef experienced mild bleaching. More *Acropora* died between six and ten weeks, but others recovered completely while more massive corals bleached. When temperatures returned to normal during the 14^th^ week, *Acropora* either died or attained complete recovery, yet, many massive colonies remained pale and started recovery. If we rely on a one-time survey without baseline data, one might have erroneously concluded species susceptibility, percentage of bleaching, and mortality in a scenario like this. By surveying bleaching at multiple time points during this event, the authors accurately captured the dynamics of bleaching and mortality and provided a foundation for future work.

Similarly, on Kavaratti Island, during the 2015 and 2016 bleaching events, the 11th-week survey found many tolerant corals mostly or entirely bleached, while thermally sensitive corals appeared fully pigmented. However, when they returned after ten months, the temperature became normal. They found a reverse pattern: susceptible corals died while resistant corals recovered in many locations. These observed bleaching patterns in two studies from sample locations are likely due to each study’s survey time difference. This contrasting conclusion underscores the importance of repetitive surveys with baseline data before taking up conservation management practices. Based on available information from the Gulf of Mannar reef, one can expect the following diagrammatic bleaching scenario in the field.

**Fig. 7.**
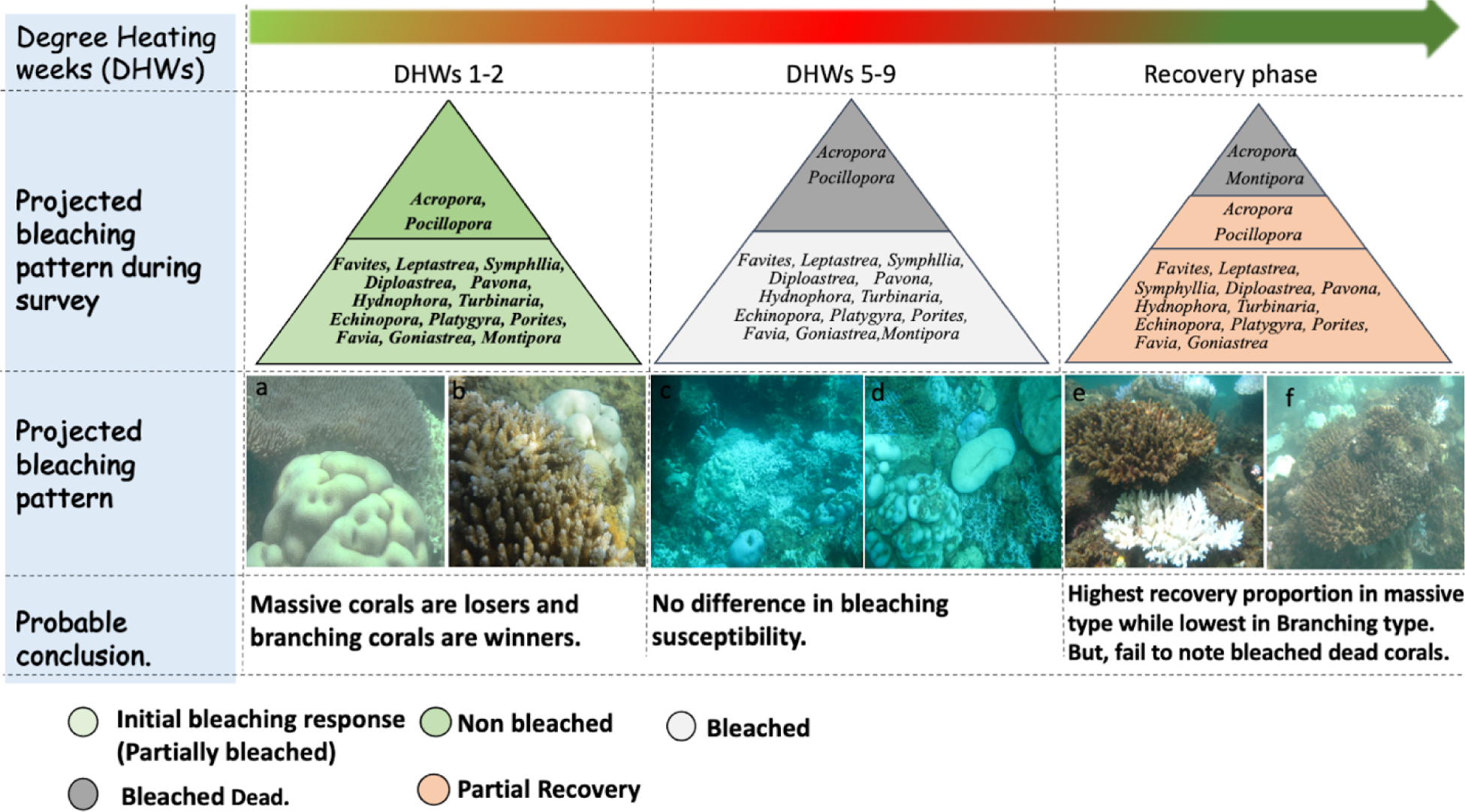
This diagram represents how a one-time survey during different DHWs could influence our result in determining corals’ susceptibility to long-term bleaching events. Probable scenarios 1) If the survey happened during DHWs 1-4, the study concludes that branching corals are more tolerant than massive corals as massive corals start to pale (partial bleaching to 10-20%) while branching corals show no bleaching/partially bleached. 2) In DHWs at peak, there is no difference in species susceptibility but most likely missed dead branching corals 3) DHWs 6-11 conclude the highest bleaching in massive corals as massive corals are being recovered. In contrast, bleached branching corals died, which divers failed to notice/record. Picture: a-b: Partially bleached Massive corals vs. unbleached branching corals; c-d: Bleached corals were irrespective of coral genera; e-f: Bleached dead *Acropora* covered with secondary algae.

## Conclusion

The frequency and severity of coral-bleaching episodes are increasing due to ongoing climate change, which alters species composition and reef function that determines future coral reefs. Hence, information on bleaching episodes, intensity, and species susceptibility is essential from broad geological location to frame conservation strategy. 23 years of concerted bleaching survey data from Indian coral reefs, i) provides insights on DHWs and their influence in bleaching intensity and species susceptibility response for major Indian reefs; ii) informed the highest susceptibility in branching corals and recovery in massive corals across locations; iv) highlighted the contrasting conclusion to long-term bleaching events and underscored the importance of repeated surveys during long-term bleaching events before concluding species susceptibility. This meta-analysis also underlines the limitations that hinder acquiring the most from our extensive analysis: i) one-time surveys with no information about DHWs, ii) no symbiont diversity data, ii) partial/incomplete bleaching information, iii) the presence and absence of data on sensitive coral genera rather than a proportion iv) lack of follow up survey to document mortality and bleaching recovery, v) contrasting in conclusions on species susceptibility to long-term bleaching events as a result of one-time survey. Despite these limitations, present study helps to improve the knowledge gap on bleaching episodes and reveal the coral community responses to short-term and long-term bleaching events. This analysis also identifies that coral bleaching studies in Indian reefs are mostly fragmented individual efforts, and based on a one-time survey. We recommend inter-institutional collaborations, adhering to a uniform survey procedure and including all details in the future survey to obtain the most accurate field data on bleaching episodes using a broad, data calibration for accuracy and quality control, and a transdisciplinary approach that will ultimately serve the common goal to improve coral health and conservation. This accurate data will not only allow us to compare with future events from Indian coral reefs and reefs worldwide, this will enable us to select potential species for restoration, which will ultimately determine restoration success and coral reef conservation.

## Acknowledgment

TT thanks the Current CSIR Pool Scientist Scheme, India. Rufford & Idea Wild small grants for support. KD is thankful to CSIR-NIO for facilities and support. Authors are thankful to Prof. Kartik Shanker, and Dr. Titus Immanuel, Centre for Ecological sciences, Indian Institute of Science for his valuable comments to improve this manuscript.

## Author contributions

**Thinesh Thangadurai**: Conceptualization, Methodology, Writing - original draft, Writing – review & editing. **Kalyan De**: Methodology, Data curation, Writing - original draft, Writing – review & editing. **Anthony Bellantuono**: Writing – review & editing. **Sobana**, **Riana Peter, Sivagurunathan:** Data curation. **Polpass Arul Jose, Joseph Selvin**: review & editing.

