## Supplementary materials.docx for "History of recurrent short and long-term coral bleaching events in Indian coral reefs: a systematic review of contrasting bleaching patterns, lessons learned, and future directions" (from another version of this preprint: version 1, 2024-05-21)

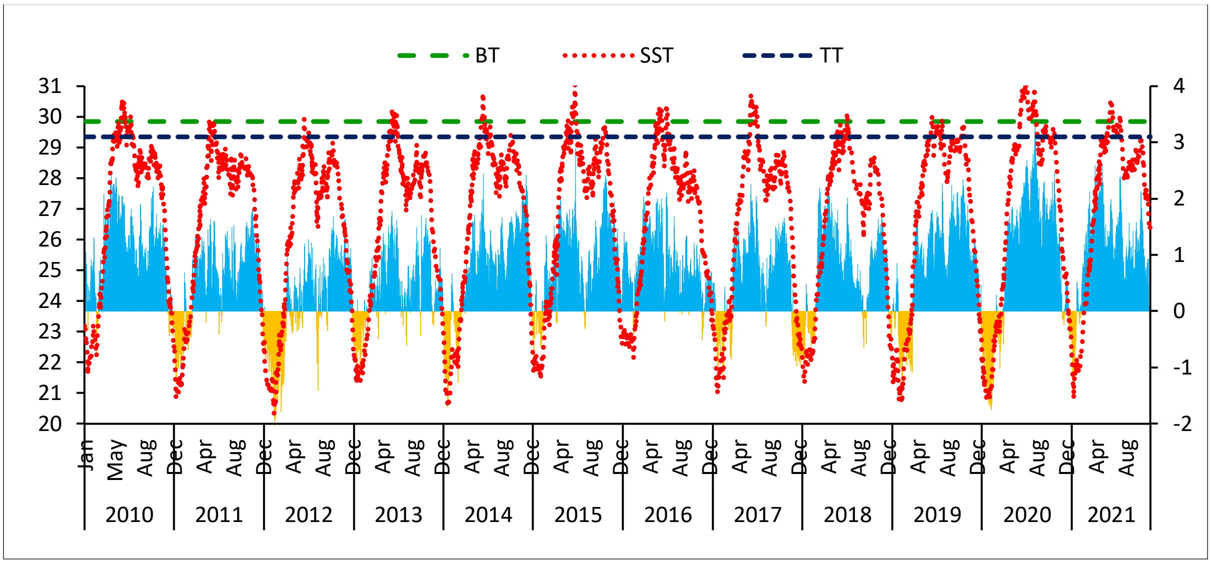


**Supplementary Fig S1:** Daily trends of mean SST and SST anomaly for the period 2010 to 2021 in Gulf of Kachchh region (Data source: NOAA CRW v 3.1). Mean daily SST and SST anomaly is given on the two 'Y' axes. The dotted red line represents the daily mean SST for the study region, while the dotted blue line shows the thermal threshold (TT), and the dotted green line represents the bleaching threshold (BT) for this same period.

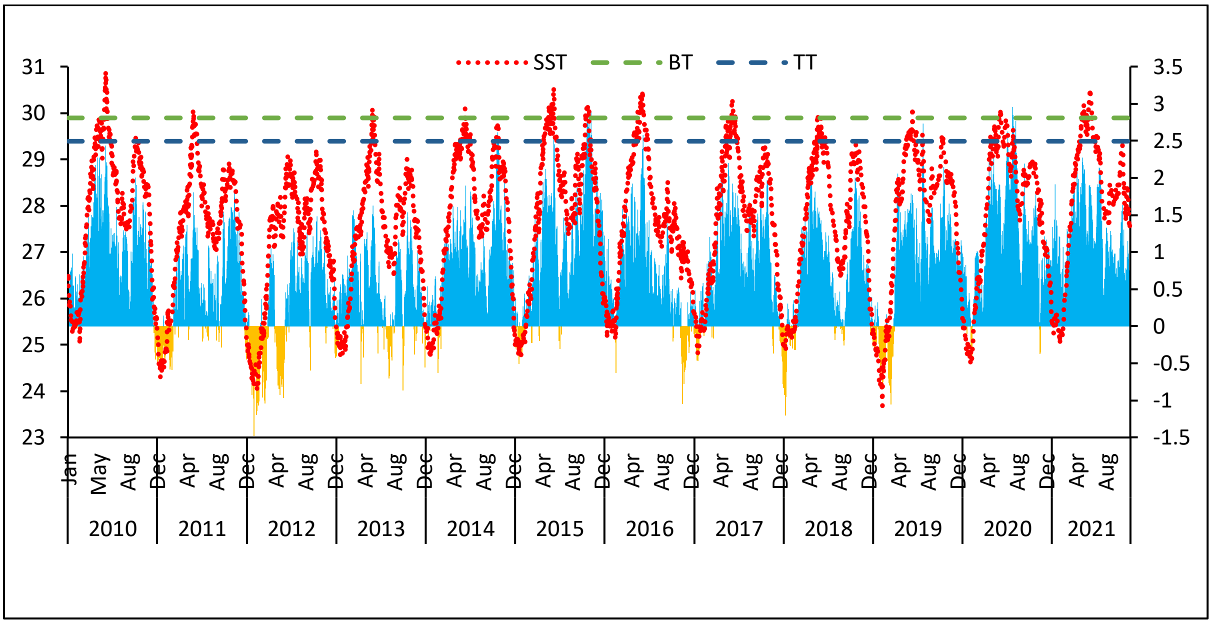

**Supplementary Fig S2:** Daily trends of mean SST and SST anomaly for the period 2010 to 2021 in west coast of India (Data source: NOAA CRW v 3.1). Mean daily SST and SST anomaly is given on the two 'Y' axes. The dotted red line represents the daily mean SST for the study region, while the dotted blue line shows the thermal threshold (TT), and the dotted green line represents the bleaching threshold (BT) for this same period.

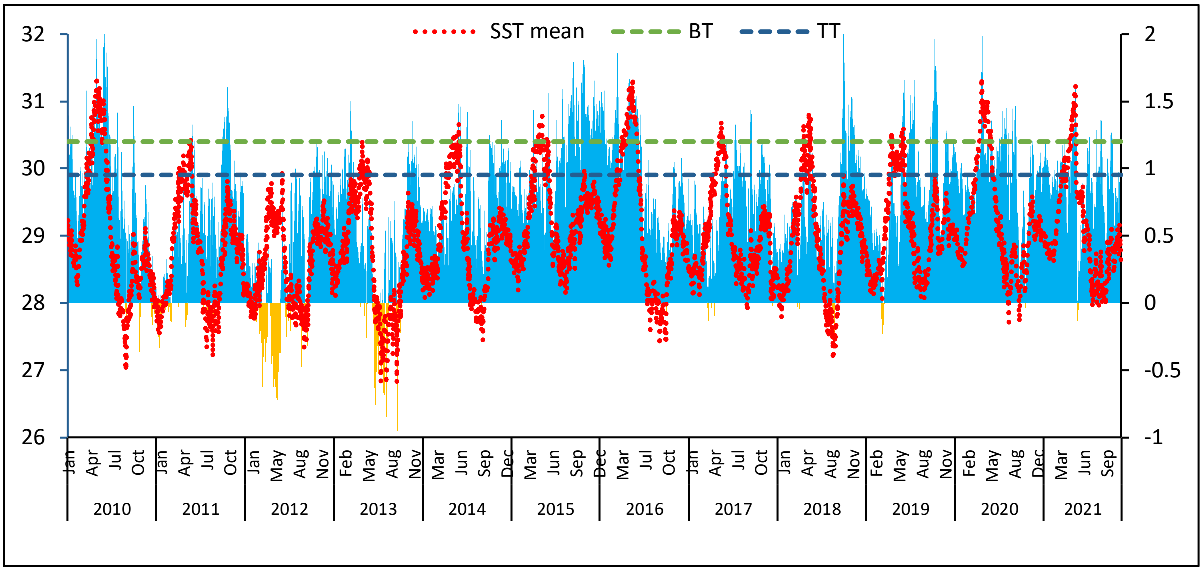


**Supplementary Fig S3:** Daily trends of mean SST and SST anomaly for the period 2010 to 2021 in Lakshadweep region (Data source: NOAA CRW v 3.1). Mean daily SST and SST anomaly is given on the two 'Y' axes. The dotted red line represents the daily mean SST for the study region, while the dotted blue line shows the thermal threshold (TT), and the dotted green line represents the bleaching threshold (BT) for this same period.

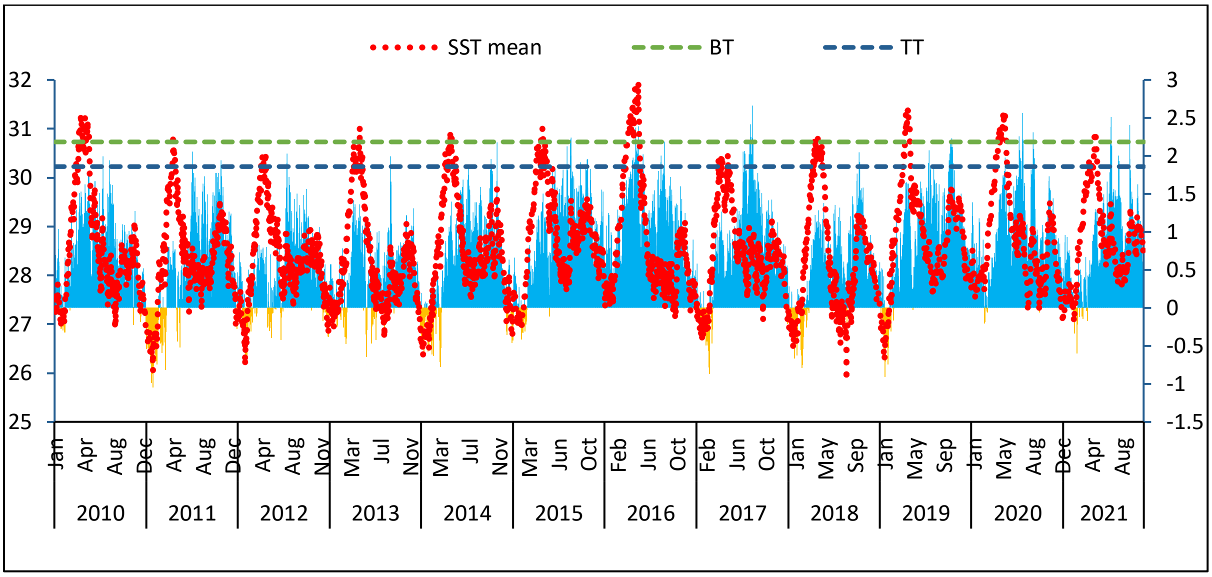


**Supplementary Fig S4:** Daily trends of mean SST and SST anomaly for the period 2010 to 2021 in Gulf of Mannar region (Data source: NOAA CRW v 3.1). Mean daily SST and SST anomaly is given on the two 'Y' axes. The dotted red line represents the daily mean SST for the study region, while the dotted blue line shows the thermal threshold (TT), and the dotted green line represents the bleaching threshold (BT) for this same period.

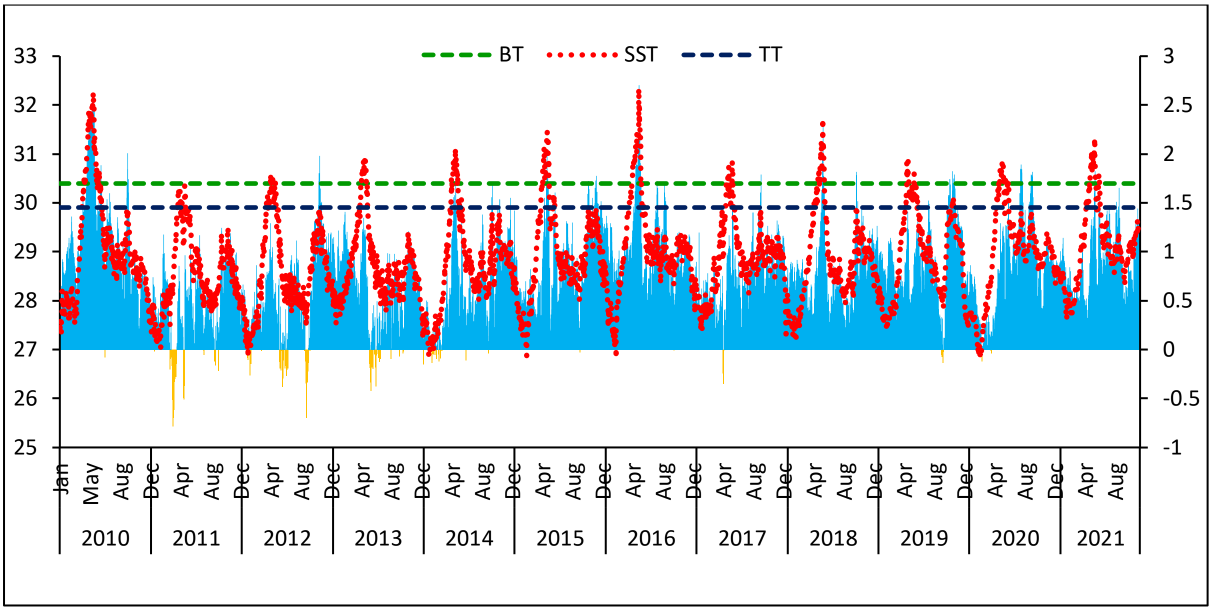


**Supplementary Fig S5:** Daily trends of mean SST and SST anomaly for the period 2010 to 2021 in Andaman region (Data source: NOAA CRW v 3.1). Mean daily SST and SST anomaly is given on the two 'Y' axes. The dotted red line represents the daily mean SST for the study region, while the dotted blue line shows the thermal threshold (TT), and the dotted green line represents the bleaching threshold (BT) for this same period.
